# The temporal recovery of contralateral and ipsilateral knee extensor torque following a bout of unilateral knee extensor resistance exercise in young, healthy resistance-trained men

**DOI:** 10.1101/2023.12.05.569582

**Authors:** Robert W. Davies, Harley L. Barnes, Brian P. Carson, Philip M. Jakeman

**Affiliations:** Chester Medical School, University of Chester, University Centre Shrewsbury, Shrewsbury, UK; Department of Physical Education and Sports Sciences, University of Limerick, Limerick, Ireland; Health Research Institute (HRI), University of Limerick, Ireland

**Keywords:** Creatine kinase, electromyography, fatigue, muscle contraction, muscle strength, quadriceps

## Abstract

The present study aimed to characterise the temporal recovery pattern of contralateral-homologous torque following a bout of unilateral resistance exercise (RE). Ten young, healthy, recreationally active, resistance-trained men performed 10 sets of 10 repetitions of knee extensor (KE) contractions at 50 % 1RM with 1 min rest between sets. Isometric maximal voluntary contraction (MVC) peak torque (PT), surface electromyography (sEMG), muscle soreness and serum creatine kinase (CK) levels were assessed immediately before and 5 min after RE cessation, and then +4 h, +24 h, +48 h and +72 h later. Data are presented as mean [95 % CI] % change from pre-exercise values. RE evoked a minor increase in CK and pain in the late recovery period (+24 h to +72 h) (P < 0.034) and decreases in ipsilateral KE PT were observed immediately post-exercise (-26 [-33, -18] %, P < 0.001) and up to +48 h (-12 [-19, -4] %, P = 0.006). Measurable decreases in PT were also observed in the non-exercised contralateral KE immediately post-exercise (-8 [-13, -3] %, P = 0.006) up to +24 h (-8 [-15, 0] %, P = 0.020), but were significantly lower than the ipsilateral KE PT (P < 0.05). These findings suggest the presence of crossover fatigue following RE in young, healthy, active, resistance-trained men, however, the magnitude and temporal recovery are substantially less severe and protracted in the contralateral homologous KE.

## Introduction

Intense unilateral muscle contractions are often accompanied by unintended muscle activity of the contralateral homologous muscle, central fatigue, and an inability to voluntarily activate (VA) both the ipsilateral and contralateral limbs (Dimitrijevic et al. 1992; Kavanagh et al. 2016). Consequently, measurable decreases occur in the torque-producing capacity of the non-exercised contralateral homologous limb, referred to as non-local muscle fatigue (NLMF) or, more specifically, crossover fatigue (CF) (Doix et al. 2013; Rattey et al. 2006; Todd et al. 2003).

NLMF has also been observed in heterologous muscles, both inferior and superior to the exercised muscle; however, the lower limbs seem particularly susceptible to NLMF/CF (Halperin et al. 2015). Conversely, there is considerable variation in the magnitude of the NLMF/CF has been reported between studies, with some failing to demonstrate any NLMF/CF whatsoever (Arora et al. 2015; Berger et al. 2010; Rattey et al. 2006), while others have (Doix et al. 2013; Halperin et al. 2014; Martin & Rattey 2007).

Similar to other non-local muscular responses to exercise (e.g., reflexes actions and cross-education), CF is thought to be mediated via interhemispheric crosstalk (Mrachacz-Kersting et al. 2018; Manca et al. 2017; Kennedy et al. 2013). Consequently, exercise bouts that involve high-intensity muscle contractions and relatively large volumes of work (e.g., resistance exercise) will result in more severe and prolonged losses of both ipsilateral and contralateral muscle torque (Halperin et al. 2015; Kawamoto et al. 2014; Hedayatpour et al. 2018; Marathamuthu et al. 2022).

Knowledge pertaining to NLMF/CF response to exercise may have implications for the delivery, planning and conduct of unilateral training/rehabilitation programmes. Thus, the purpose of the present study was to characterise the temporal recovery pattern of contralateral and ipsilateral knee extensor (KE) torque following a bout of unilateral KE resistance exercise (RE). In the present study, both ipsilateral and contralateral KE muscle torque, surface electromyography (sEMG) and muscle soreness were measured immediately before and after RE, four hours later, and then at 24 h intervals for the next 3 days in a cohort recreationally active participants who regularly participate in RE.

## Methods

### Participants and Ethics

The study design and all procedures were approved by the University of Limerick Education and Health Sciences Research Ethics Committee (2014_05_11_EHS), conforming to standards set by the Declaration of Helsinki. Participants were informed of the risks and benefits associated with participation before providing written consent. Eligibility criteria were: (i) young adults aged 18 to 35 y; (ii) resistance trained and strong (defined as > 0.5 y resistance training experience > 3 h per wk and an isometric MVC PT > 2.5 N·m·kg^-1^ body mass) (iii) in good health with no injury, illness or history of cardiovascular, metabolic, neurological or motor disorders; (iv) not currently taking any medication. To assess eligibility criteria, participants reported to the laboratory on two separate days the week before testing for a health screening, familiarisation testing and MVC torque tests. Ten participants completed the study in full (age 23 (8) y; stature 1.80 (0.07) m; body mass 74.2 (7.4) kg; 2.2 (0.8) y resistance training experience; mean (SD)).

### Study Protocol

Participants reported to the laboratory in the morning (06.00 to 10.00 am) for five consecutive days, at the same time each day. Participants fasted overnight (∼ 10 h postabsorptive) and consumed only 500 ml of water in the hour before testing. Pre-exercise knee extensor MVC torque, sEMG and pain responses were assessed in both the ipsilateral (i.e., exercised) and contralateral (i.e., non-exercised) knee extensors. Participants then performed 200 intermittent unilateral knee extensions with their dominant limb on an isokinetic dynamometer (Contrex, Dubendorf). Continuous concentric and eccentric contractions were alternated using real-time visual feedback. For each contraction, participants were required to meet a torque target (50 % PT). Knee extension dynamic range was 90° (90° to 180º knee angle) at 90°·s^-1^. Every 20 contractions, 1 min rest was taken (0.25 duty cycle). Total work during the exercise session was 24 (4) kJ, equally divided between the eccentric (12 (2) kJ) and concentric (12 (1) kJ) contractions. Measures were then repeated 5 min post-exercise, +4 h, +24 h, +48 h, and +72 h post-exercise (Figure 1).

**Figure 1.**
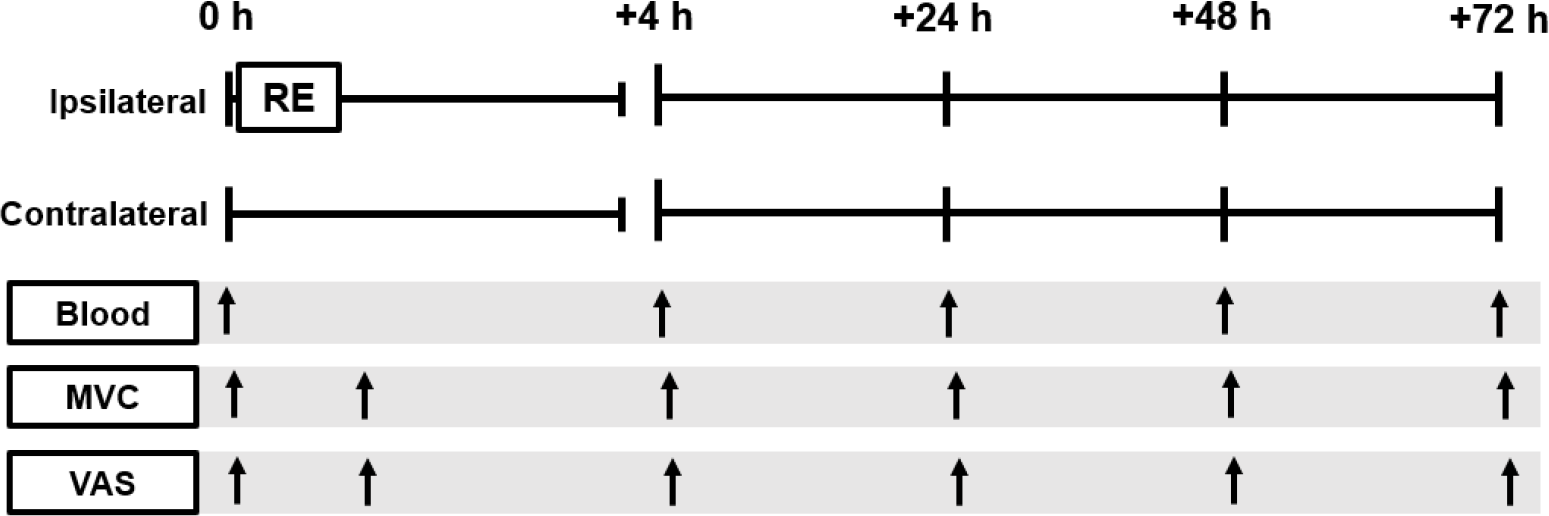
Schematic diagram of the study protocol. MVC, Maximal voluntary contraction; VAS, Visual analogue scale measures; RE, Unilateral Resistance Exercise.

### Muscle Torque Assessment

An isokinetic dynamometer (Contrex, Dubendorf) was used to measure isometric MVC torque. Participants were seated at a 90º hip angle with the rotational axis of the dynamometer aligned with the lateral femoral epicondyle, the shin pad of the lever arm attached 3 cm proximal to the lateral malleolus and strapping was applied to the chest, pelvis and mid-thigh to isolate the knee extensor action. Three 5 s isometric MVCs were performed 60º below full extension with 3 min rest between each contraction. Participants were instructed to push as fast and forcefully as possible. Standard verbal encouragement was provided during the MVC. The dynamometer strain-gauge signal was sampled at 256 Hz. Isometric PT was defined as the highest value observed during the three attempts. The coefficient of variation (CV) from the three repeated baseline MVCs was 4.7 %.

### Electromyography

sEMG was measured during all MVCs. Prior to measurement, skin was prepared (shaving, abrasion, and cleaning with ethanol), and circular pairs of surface electrodes (30mm inter-electrode distance, Ag/AgCl UTP 3.2 mm) were placed on the belly of the *m. vastus lateralis, m. vastus medialis* and *m. rectus femoris* following SENIAM guidelines (Hermans et al. 2000). Electrodes were contoured with an indelible marker before removal to ensure the same position for each test. Electrodes were connected directly to differential signal amplifiers and digitised to a computer interface receiver at 1000 Hz (MyoSystem, Noraxon, AZ, USA). Data were digitally high-pass filtered through a 1^st^ order zero-lag Butterworth filter with a 5 Hz cut-off followed by a root-mean-square with a 50 ms time constant (MyoResearch, Noraxon, AZ, USA). For this study, the relationship between knee extensor torque and the overall sEMG pattern was more important than the behaviour of the individual muscles. Therefore, sEMG data from the individual muscles was summed prior to analysis (Viitasalo, 1984). Peak amplitude was the highest voltage, and iEMG_max_ was the highest 1 s integrated sEMG (∫ V d*t*) during the MVC.

### Serum Creatine Kinase

Venous blood samples were taken from the antecubital fossa, pre-exercise, 5 min post-exercise, +4 h, + 24 h, +48 h, and +72 h post-exercise. Whole blood samples were clotted at room temperature and then centrifuged at 3500 RPM for 10 min at 20°C. Serum was separated from whole blood and stored at -80°c prior to analysis. Analysis of the muscle damage marker creatine kinase (CK) was quantified using spectrophotometry using a coupled enzyme-linked reaction assay (Sigma-Aldrich). Samples were measured in duplicate. The intra-assay coefficient of variation (CV) for quintuplicate samples was < 3%, inter-assay CV was < 10%.

### Muscle Pain Responses

Before the MVCs, the pressure-induced pain response was evoked by a custom-made algometer. A cylindrical plastic probe with a 10 mm flat head was incrementally applied to evoke a mild pressure-pain response for each muscle at the belly of the muscle. (sEMG electrode placement sites). Participants verbally confirmed minimum pressure-pain response. The force applied was then recorded (∼ 50 N) and repeated for each subsequent measurement. The average pain response across the three sites was analysed. Participants were also asked to evaluate muscle pain and soreness during the MVC (i.e., contraction-induced pain/soreness). Participants indicated pain responses by marking a 100mm visual analogue scale (VAS). VAS descriptors ranged from ‘not at all’ (0 mm) to ‘extremely’ (100 mm).

### Statistical Analysis

Data were assessed for normality, sphericity, and homogeneity of variance prior to statistical analysis (no violations were observed). Two-way (2×6) repeated measures analysis of variance (ANOVA) was used to determine limb × time interaction. Repeated measures ANOVA was used to assess time effects for each limb and CK. Post-hoc t-tests were used to assess effects over time and between limbs. The critical significance level was α = 0.05. Data are reported as mean (SD) or mean percent change [95% CI]. The magnitude of the change was examined by effect size calculation (d) (Lakens, 2013). Participant responses were also examined individually using their baseline (pre-exercise) MVC PT typical error (TE). Post-exercise MVC PTs 1.96 TEs below baseline were deemed as a genuine loss of PT (Swinson et al. 2018). Statistical analysis was performed in Jamovi (2.2.5).

## Results

### Knee Extensor Torque and sEMG

There was a limb × time interaction (P < 0.001) with time main effects for both limbs (P < 0.001). Substantial decreases in PT were observed 5 min post-exercise (-26 [-33, -18] %, d = 2.2, P < 0.001), +4 h (-18 [-23, -12] %, d = 2.0, P < 0.001), +24 h (-16 [-23, -9] %, d = 1.3, P = 0.001), +48 h (-12 [-19, -4] %, d = 1.0, P = 0.006) (Table 1, Figure 2). Loss of PT was observed for all 10 participants 5 min post-exercise, 9 participants at +4 h, 7 participants at +24 h, and 6 participants at +48 h post-exercise. Recovery of PT in the ipsilateral limb was observed +72 h post-exercise (0 [-7, 9] %, d = 0.0, P = 0.478) with 8 out of 10 participants returning to baseline values.

**Table 1.** Change in knee extensor (KE) in maximal voluntary contraction (MVC) peak torque (PT) and pressure-pain response (PPR) of the exercised (ipsilateral) and non-exercised (contralateral) KE following unilateral resistance exercise. Data are mean (SD). Post, 5 min post-exercise; CK, serum creatine kinase concentration. *P < 0.1, **P < 0.05, ***P < 0.01 different from pre-exercise (Pre). ^**†**^ limb × time (Pre) (P < 0.05)

|  | Pre | Post | +4 h | +24 h | +48 h | +72 h |
| --- | --- | --- | --- | --- | --- | --- |
| **PT (N·m)** |  |  |  |  |  |  |
| Ipsilateral | 229 (26) | 173 (34)*** | 189 (32)*** | 193 (36)*** | 202 (35)*** | 229 (40) |
| Contralateral | 220 (32) | 203 (30)***† | 207 (25)*† | 202 (28)**† | 212 (31)† | 221 (31) |
| **PPR (mm)** |  |  |  |  |  |  |
| Ipsilateral | 1.4 (1.4) | 1.6 (1.3) | 1.9 (1.6) | 2.5 (1.9)*** | 2.8 (2.1)** | 2.5 (1.8)** |
| Contralateral | 1.6 (1.2) | 1.6 (1.3) | 1.9 (1.6) | 2.1 (1.8)† | 1.9 (1.5)† | 2.2 (1.9) |
| **CK (IU·L<sup>-1</sup>)** |  |  |  |  |  |  |
|  | 98 (61) | 99 (55) | 130 (52)*** | 153 (47)*** | 143 (66)** | 162 (85)** |

**Figure 2.**
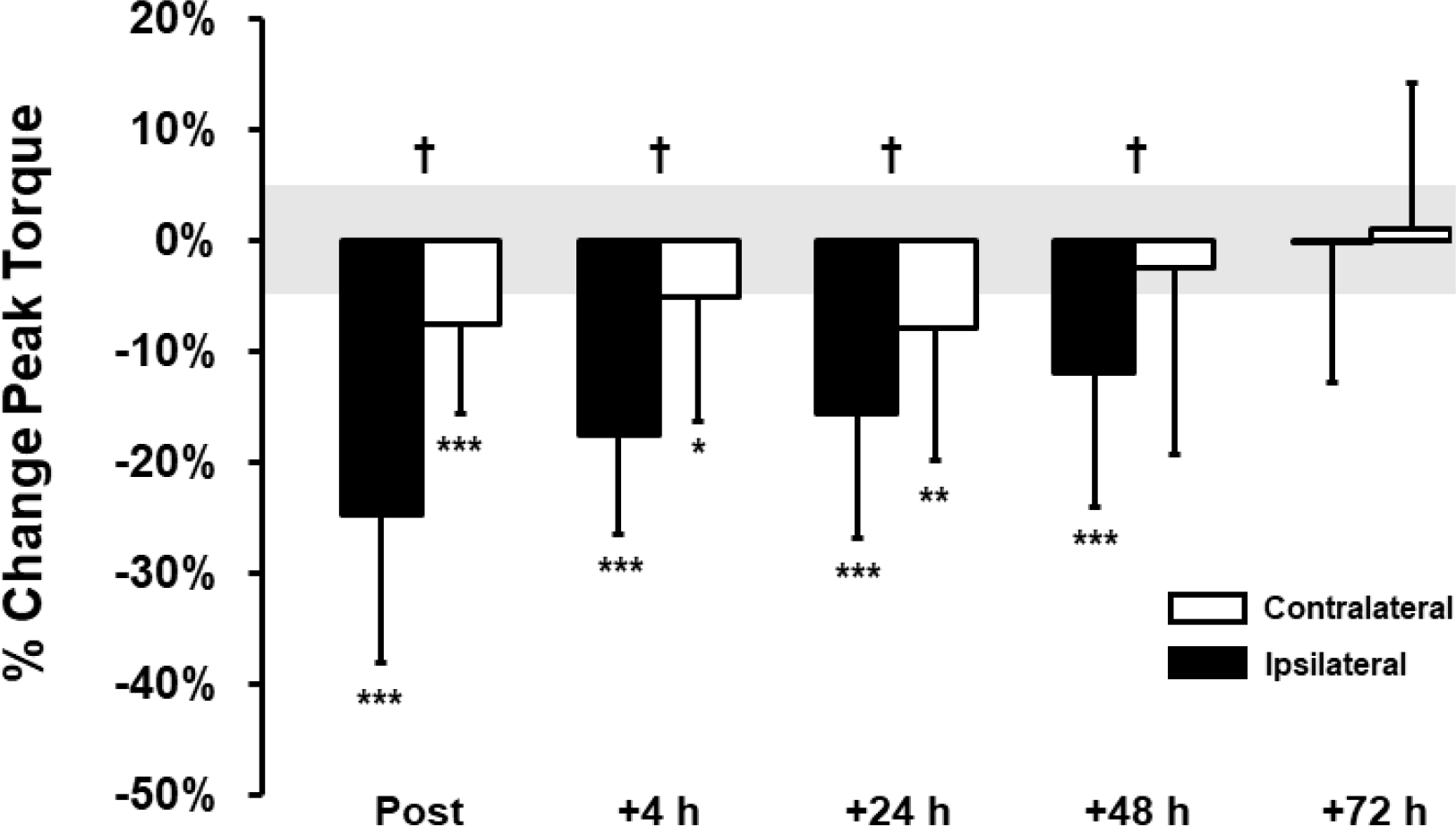
Knee extensor (KE) isometric maximal voluntary contraction peak torque, percent change from pre-exercise. Data are mean (SD) for non-exercised contralateral KE (open bars) and exercised ipsilateral KE (black bars) following unilateral KE resistance exercise bout. Post, 5 min post-exercise. ^**†**^ difference between limbs (P < 0.05); different to pre-exercise *P < 0.10, **P < 0.05, ***P < 0.01. Grey shaded area is the coefficient of variation (CV) pre-exercise (4.7 %).

Whilst substantially lower than the ipsilateral limb (P < 0.05), measurable decreases in PT were observed 5 min post-exercise in the contralateral limb (-8 [-13, -3] %, d = 1.0, P = 0.006). Moreover, there were also measurable decrements in contralateral PT +4 h (-5 [-12, 2] d = 0.5, P = 0.066) and +24 h (-8 [-15, 0] %, d = 0.8, P = 0.020) post-exercise (Table 1, Figure 2). Recovery of PT was observed in the contralateral limb, returning to pre-exercise levels by +48 h (-3 [-13, 8] %, d= 0.2, P = 0.235) and +72 h (1 [-7, 9] %, d = 0.0, P = 0.529) post-exercise. Loss of PT in the contralateral KE was observed in 6 participants at 5 min, 5 participants at +4 h, +24 h and +48 h and 2 participants at +72 h post-exercise.

Although a moderate positive correlation was observed between post-exercise contralateral and ipsilateral torque (r = 0.4, P = 0.007), there was substantial variation between participants. Indeed, CF was observed in only 63 % of the instances where measurable loss of ipsilateral PT occurred. Moreover, no CF was observed in 3 participants despite presenting with significant ipsilateral PT decrements. We report no interaction or time main effect in any sEMG parameter in either limb (P > 0.434).

### Muscle Pain and Muscle Damage

There was a limb × time interaction for the pressure-pain response (P = 0.034) but not for the contraction pain response (P = 0.664). A time main effect of the pressure-pain response was only observed in the ipsilateral limb (P = 0.009), where delayed increases were observed +24 h (+1.1 [0.4, 1.8] cm, d = 1.0, P = 0.007) and +48 h (+ 1.3 [0.4, 2.3] cm, d = 0.9, P = 0.013) and + 72 h (+1.0 [0.1, 1.9] cm, d = 0.7, P = 0.032) post-exercise. There was no change in the pain response in the contralateral limb over time whatsoever (P = 0.598) (Table 1).

An increase in serum CK was observed over time (P = 0.009). Increases from pre-exercise values (98 (61) IU·L^-1^) were observed +4 h (+31 [13, 50] IU·L^-1^, d = 1.2, P = 0.002), +24 h (+54 [18, 96] IU·L^-1^, d = 0.9, P = 0.008), +48 h (+45 [6, 96] IU·L^-1^, d = 0.6, P = 0.041) and +72 h (+63 [4, 130] IU·L^-1^, d = 0.7, P = 0.032) (Table 1).

## Discussion

The present study aimed to characterise the temporal recovery pattern of contralateral and ipsilateral KE PT following a bout of unilateral RE in a cohort of young, healthy, and active men. Post-RE, decrements in ipsilateral PT were observed immediately and persisted up to +48 h, while decreases in PT of the contralateral homologous non-exercised KE persisted up to +24 h. However, the loss of PT in the contralateral KE was substantially lower than in the ipsilateral KE (17% *vs* 7% average loss of PT over the first 24 h post-exercise), with one-third of the participants exhibiting no CF whatsoever. Nevertheless, deficits in PT were significantly correlated between exercised and non-exercised limbs, suggesting the magnitude and the time course of recovery of PT in the homologous contralateral and ipsilateral muscles were related.

In the extant literature, CF has conventionally been examined during or immediately after exercise (Amann et al. 2013; Kawamoto et al. 2014). To our knowledge, this is the first study to characterise the full temporal recovery of both contralateral and ipsilateral homologous KE, reporting mild/moderate but measurable and long-lasting CF up to +24 h post-exercise, and possibly longer in some individuals. While two other studies have previously reported long-lasting CF up to +72 (Hedayatpour et al. 2018; Marathamuthu et al. 2022), these studies utilised novel forms of (eccentric) exercise designed to purposefully evoke ultrastructural damage, oedema, muscle pain and stiffness in the exercised muscle – resulting in a severe and prolonged decrease in MVC torque (Allen et al. 2008; Clarkson & Hubal, 2002; Halperin et al. 2014).

Conversely, in the present study, we aimed to characterise the NLMF/CF response to an ecologically valid mode of RE (i.e., submaximal, familiar, and isotonic) in participants who are regularly partaking in RE. Consequently, it was not our intent to purposefully induce large levels of muscle damage, muscle soreness, or loss of muscle function via experimental modes of exercise (e.g., maximal/supramaximal, unfamiliar, eccentrically biased), which are not typically undertaken in normal training or everyday life. Thus, the level of muscle damage, pain/soreness, and loss of PT in the ipsilateral limb in the present study was considerably lower than after purposefully damaging experimental modes of eccentric exercise (Hedayatpour et al. 2018; Marathamuthu et al. 2022). Consequently, this may also explain the truncated and less severe levels of CF, and the absence of any measurable change in any sEMG parameter.

Indeed, some studies have reported commensurate decrements in sEMG amplitude alongside NLFM/CF (Hedayatpour et al. 2018; Rattey et al. 2006), but others have not (Aboodarda et al. 2015; Halperin et al. 2014; Kawamoto et al. 2014; Šambaher et al. 2016). Although there is an established relationship between voluntary sEMG and torque output in the larger lower-body muscle groups, the relationship is non-linear (i.e., decreases in muscle torque can occur in the absence of any change in the EMG signal) (Kooistra et al. 2007; Lippold, 1952). We suspect the absence of any change in sEMG in the present study may be related to the relatively low decrements in muscle torque; caused by physiological alterations that are not reflected/measurable via evaluation of the MVC-sEMG (e.g., change in firing rate, supraspinal and/or corticospinal excitability) (Kawamoto et al. 2014). Further research is needed to determine the underlying mechanisms causing the low-level CF following this mode of exercise.

Data from the extant literature suggest that the lower-body muscle groups (i.e., predominately quadriceps) are more susceptible to CF than upper-body muscles (e.g., biceps brachii and dorsal interossei muscles) (Halperin et al. 2015). Moreover, it has also been suggested that training experience, sex, and the type of outcome measure also moderate NLMF/CF effects (Halperin et al. 2015). Therefore, we emphasise that the findings from this study are limited MVC PT output of the KE muscles in young, healthy resistance-trained men. Additionally, despite a relatively homologous group of participants, there was substantial unexplained variation in the magnitude and temporal pattern of recovery of both KEs. Indeed, no measurable CF was observed in three participants whatsoever despite presenting with significant loss of PT in the ipsilateral KE.

In conclusion, following a unilateral bout of KE RE, moderate levels of CF were observed in the contralateral homologous KE, which recovered by +48 h. These changes occurred alongside relatively mild/moderate increases in exercise-induced muscle damage and muscle pain/soreness and moderate/large deficits PT in the ipsilateral KE, sustained up to +48 to 72 h post-exercise. These findings should be taken into consideration in the development of unilateral RE or rehabilitation programmes as low-level but measurable residual CF occurs following RE.

## Acknowledgements

The authors wish to acknowledge the participants for their time and Paul Nevin for his technical expertise.

## References

Aboodarda SJ, Šambaher N, Behm DG. Unilateral elbow flexion fatigue modulates corticospinal responsiveness in non‐fatigued contralateral biceps brachii. Scand J Med Sci Sports 2016;26(11):1301–1312.

Allen DG, Lamb G, Westerblad H. Skeletal muscle fatigue: cellular mechanisms. Physiol Rev 2008;88:287–332.

Amann M, Venturelli M, Ives SJ, McDaniel J, Layec G, Rossman MJ, Richardson RS. Peripheral fatigue limits endurance exercise via a sensory feedback-mediated reduction in spinal motoneuronal output. J Appl Physiol 2013;115(3):355–364.

Arora S, Budden S, Byrne JM, Behm DG. Effect of unilateral knee extensor fatigue on force and balance of the contralateral limb. Eur J Appl Physiol 2015;115: 2177–2187.

Berger LL, Regueme SC, Forestier N. Unilateral lower limb muscle fatigue induces bilateral effects on undisturbed stance and muscle EMG activities. J Electromyogr Kinesiology 2010;20:947–952.

Clarkson PM, Sayers SP. Etiology of exercise-induced muscle damage. Can J Appl Physiol 1999; 24:234–248.

Doix AC, Lefevre F, Colson SS. Time course of the crossover effect of fatigue on the contralateral muscle after unilateral exercise. PLoS One 2013;8:e64910.

Dimitrijevic MR, McKay WB, Sarjanovic I, Sherwood AM, Svirtlih L, Vrbova G. Coactivation of ipsi- and contralateral muscle groups during contraction of ankle dorsiflexors. J Neurol Sci 1992;109:49–55.

Halperin I, Aboodarda SJ, Behm DG. Knee extension fatigue attenuates repeated force production of the elbow flexors. Eur J Sport Sci 2014;14:823–829

Halperin I, Chapman DW, Behm DG. Non-local muscle fatigue: effects and possible mechanisms. Eur J Appl Physiol 2015;115(10):2031–2048.

Hermens HJ, Freriks B, Disselhorst-Klug C, Rau G. Development of recommendations for SEMG sensors and sensor placement procedures. J Electromyogr Kinesiol 2000;10(5):361–374.

Hedayatpour N, Izanloo Z, Falla D. Effects of eccentric exercise and delayed onset of muscle soreness on the homologous muscle of the contralateral limb. J Electromyogr Kinesiol 2018;41:154–159.

Kavanagh JJ, Feldman MR, Simmonds MJ. Maximal intermittent contractions of the first dorsal interosseous inhibits voluntary activation of the contralateral homologous muscle. J Neurophysiol 2016;116(5):2272–80.

Kawamoto JE, Aboodarda SJ, Behm DG. Effect of differing intensities of fatiguing dynamic contractions on contralateral homologous muscle performance. J Sports Sci Med 2014;13(4):836–845.

Kennedy A, Hug F, Sveistrup H, Guevel A. Fatiguing handgrip exercise alters maximal force-generating capacity of plantar-flexors. Eur J Appl Physiol 2013; 113:559–566.

Kooistra RD, de Ruiter CJ, de Haan A. Conventionally assessed voluntary activation does not represent relative voluntary torque production. Eur J Appl Physiol 2007;100(3):309–320.

Lippold, OCJ. The relation between integrated action potentials in a human muscle and its isometric tension. J Physiol 1952;117(4):492–499.

Manca A, Dragone D, Dvir Z, Deriu F. Cross-education of muscular strength following unilateral resistance training: a meta-analysis. Eur J Appl Physiol 2017;117(11):2335–2354.

Marathamuthu S, Selvanayagam VS, Yusof A. Contralateral Effects of Eccentric Exercise and DOMS of the Plantar Flexors: Evidence of Central Involvement. Res Q Exerc Sport. 2022 Jun;93(2):240–249.

Martin PG, Rattey J. Central fatigue explains sex differences in muscle fatigue and contralateral crossover effects of maximal contractions. Pflügers Arch 2007;454:957–969.

Mrachacz-Kersting N, Gervasio S, Marchand-Pauvert V. Evidence for a supraspinal contribution to the human crossed reflex response during human walking. Front Human Neurosci 2018;12:260.

Rattey J, Martin PG, Kay D, Cannon J, Marino FE. Contralateral muscle fatigue in human quadriceps muscle: evidence for a centrally mediated fatigue response and crossover effect. Pflügers Arch 2006;452:199–207.

Šambaher N, Aboodarda SJ, Behm, DG. Bilateral knee extensor fatigue modulates force and responsiveness of the corticospinal pathway in the non-fatigued, dominant elbow flexors. Front Hum Neurosci 2016;10:18.

Swinton PA, Hemingway BS, Saunders B, Gualano B, Dolan E. A statistical framework to interpret individual response to intervention: paving the way for personalized nutrition and exercise prescription. Front Nutr. 2018;28:541.

Todd G, Petersen NT, Taylor JL, Gandevia SC. The effect of a contralateral contraction on maximal voluntary activation and central fatigue in elbow flexor muscles. Exp Brain Res 2003;150(3):308–313.

Viitasalo JT. Electromechanical behaviour of the knee extensor musculature in maximal isometric and concentric contractions and in jumping. Electromyogr. Clin. Neurophysiol. 1984;24(4):293–303.

